# Synergies on many levels: Microbially driven redoxgradients as drivers of greenhouse gas emissions in a pH neutral fen

**DOI:** 10.1101/2025.11.10.687636

**Authors:** Katharina Kujala, Viktoriia Berezovska, Johannes Cunow, Sherif Hegazy, Elisa Heilmann, Marcus A. Horn, Iida Höyhtyä, Petra Korhonen, Essi Lyytinen, Jennyffer Martinez Quimbayo, Francisca Prieto-Fernández, Lassi Päkkilä, Veera Pöykiö, Sara Solastie, Laura Tarvainen

## Abstract

Pristine northern peatlands store large amounts of carbon and nitrogen which are only slowly released under water-logged conditions due to dominance of anaerobic degradation processes. While different parts of the system such as GHG emissions have been intensively studied in many places, less focus has been paid to the integration of different peat processes. A field course at Puukkosuo fen, a pH-neutral site in northern Finland, aimed to investigate the versatile microbial communities in Puukkosuo fen, assess their metabolic potential and interpret *in situ* peat and porewater observations based on the context of the detected microbial communities and process potential. Microbial communities harbored microorganisms with aerobic and anaerobic metabolic capabilities for nitrification, denitrification, iron reduction, sulfate reduction, fermentation and methanogenesis. Most probable number counts demonstrated that groups like aerobic and anaerobic heterotrophs, denitrifiers and sulfate reducers where highly abundant, while microcosm incubations highlighted the process potential of different functional groups, especially in surface peat. Measurements of *in situ* concentrations of process substrates, intermediates or products in peat and porewater indicated that different anaerobic processes were active in anoxic peat layers which synergistically contributed to organic matter degradation in Puukkosuo fen.

## Introduction

The peatlands of the northern hemisphere constitute one of the biggest terrestrial carbon and nitrogen reservoirs (Gorham 1991; Yu 2012). Peat forms in areas where water supply exceeds drainage and evapotransporation, leading to soil saturation, e.g., due to low soil hydraulic conductivity and rather flat topography. Peat soil consists of poorly decomposed parts of plants which accumulate in water-logged conditions. The water table often stays close to the soil surface and the deeper layers of peat remain anoxic due to limited diffusion of oxygen. Peatlands have a large impact on greenhouse gas cycling: in general, pristine peatlands are net sinks of carbon dioxide (CO_2_) and net sources of methane (CH_4_), and they can be a minor source or sink of nitrous oxide (N_2_O) depending on the conditions (Takakai *et al*. 2008; Kolb and Horn 2012; Qiu *et al*. 2022).

In subsurface peat where light and oxygen are not available, organic matter decomposition rates are low and governed by microorganisms capable of fermentative processes or of anaerobic respirations using alternative terminal electron acceptors such as nitrate, iron (Fe), sulfate, or CO_2_ (Todorova, Siegel and Costello 2005; Drake, Horn and Wüst 2009). Initial steps in the degradation of organic matter include hydrolysis of polymers such as cellulose, lignin or chitin to monomers. Monomers are then further metabolized by primary or secondary fermenters (producing short-chain fatty acids, alcohols, H_2_ and CO_2_). Fermentation products as well as monomers serve as electron donors for many anaerobic respirations such as denitrification, Fe or sulfate reduction, as well as acetogenesis (Drake, Horn and Wüst 2009). When no other terminal electron acceptors are available, CO_2_ is reduced to methane or acetate by methanogens or acetogens, respectively (Drake, Horn and Wüst 2009).

While theoretically, terminal electron acceptors would be used sequentially depending on the energy yield, different types of anaerobic respiration often occur in parallel (in the same site, at the same time), and the contribution of each process is governed by a multitude of factors including electron donor and acceptor concentrations, the abundance of microbial functional groups, substrate affinities of enzymes as well as environmental parameters like temperature or pH. Microbial communities in peatlands are not only shaped by adaptation of the metabolic capabilities of its members to the available resources and environmental conditions, but also by competition for resources on one hand as well as synergistic interactions on the other hand. Thus, to understand organic matter degradation in water-logged peat soil, considering single microbial groups or monitoring single processes is insufficient to grasp the complete picture.

The presented study is the outcome of a course organized for MSc-level and doctoral students at the Oulanka research station of the University of Oulu. Just like the peat microbes working together to degrade organic matter in peat, the course participants worked together to probe different processes and parts of the peat microbial community, which in the end were integrated to provide a more complete representation of peat microbial processes. The participants were divided into five groups of two to three and were assigned a functional group on which to focus during the course (namely, nitrifiers, denitrifiers, Fe(III) reducers, sulfate reducers or fermenters and methanogens). Participants set up incubations and MPN counts targeting their assigned physiological group and contributed to group efforts such as sampling, analysis of peat and porewater as well as analysis of microbial community composition. After the course, participants calculated results and reported their findings in a “mini-publication”. The here presented work is a synthesis stemming directly from these “mini-publications”, and the course has thus fulfilled a double purpose in education and generation of valuable scientific data.

## Material and Methods

### Study site and sampling procedures

The study was conducted in Puukkosuo, a pH-neutral fen located in northeastern Finland. Puukkosuo is a meso-eutrophic fen with a plant cover consisting mainly of mosses and sedges. Mean annual temperature and precipitation at the site are 1° C and 556 mm (average for years 2001-2025, based on data from the Finnish Meteorological Institute), with July being the warmest month (on average 15.7° C) and January the coldest (on average -13.0° C). The site is snow-covered from October to May. Peat and porewater samples were collected in September 2023. Peat samples were collected on two occasions using a Russian soil corer. On September 14^th^, samples were obtained from 0 to 20 cm and 40 to 60 cm depth. These samples were homogenized, stored at 4° C for five days and used for MPN counts and microcosm incubations. On September 19^th^, 2023, three replicate peat cores spanning depths from 0 to 80 cm below the peat surface were obtained (Fig. 1). Cores were partitioned into 10 cm sections which were homogenized on site, and subsamples were stored at -20° C for later DNA extraction and 4° C for all other analyses upon arrival in the laboratory (< 2 h after sampling). For collection of porewater samples, six equilibrium dialysis chambers (peepers) with a depth-resolution of 1 cm as described in (Eberle *et al*. 2021), were installed in the fen and left to equilibrate for approximately 4 weeks. Peepers were retrieved on September 19^th^, 2023, sealed with rubber plates and wrap to avoid oxygen exposure and transported to the laboratory for immediate further treatment.

**Figure 1:**
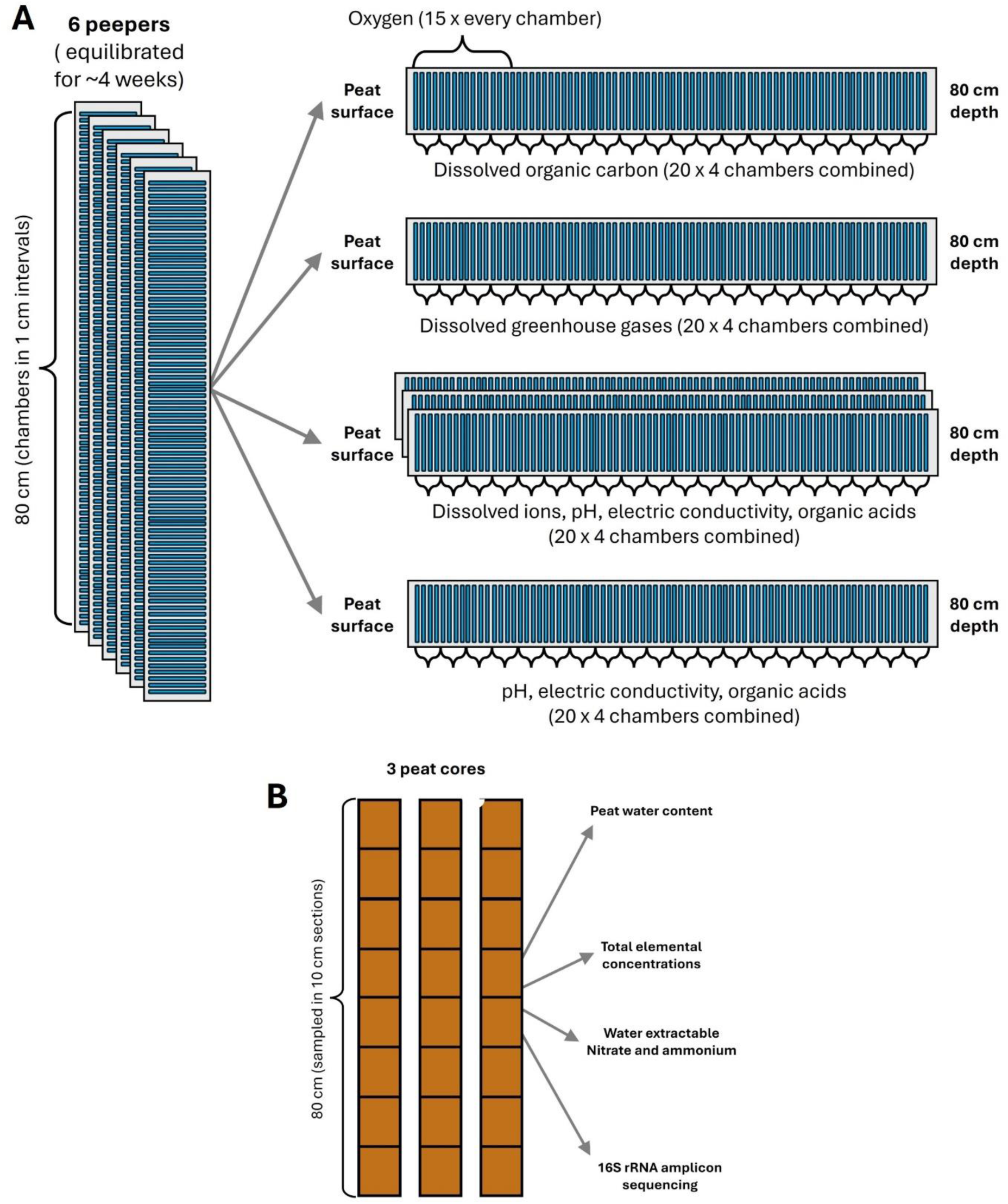
Schematic overview of field sampling and sample analysis. A) Six porewater samplers were installed in the fen on August 24th, 2023, and retrieved on September 19th, 2023. Water from the porewater samplers was then subjected to the indicated analyses. B) Three peat cores were taken on September 19^th^, 2023. Cores were divided into 10 cm sections, peat of each section was homogenized and subjected to the indicated analyses.

### Porewater analyses

The six peepers retrieved from the fen were used for different analyses (Fig. 1). For the determination of dissolved oxygen, peeper chambers were uncovered one by one to avoid oxygen penetration into the chambers. The peeper membrane was pierced with a needle, and a microsensor (Unisense OX-MR, Unisense, Denmark) was inserted halfway into the underlying chamber for measurement of oxygen concentration. Measurements were recorded for approximately 30 sec before moving to the next chamber. Dissolved oxygen was only measured in the top 15 chambers, as the concentration dropped to 0 already at 11 cm from the top.

For all other measurements, water from four adjacent chambers was pooled to obtain sufficient sample volume for the analyses, yielding a total of 20 samples per peeper (4 cm depth resolution). Samples were extracted using a syringe and needle. One peeper was used for the determination of dissolved greenhouse gases (CO_2_, N_2_O, CH_4_). In a syringe, ∼20 mL total sample volume (exact volume noted) was mixed with 30 mL of room air. The syringe was closed with a three-way-valve and vigorously shaken for 3 minutes to allow for the formation of an equilibrium between the porewater and the syringe headspace. 20 mL of the headspace gas were transferred into pre-evacuated 15 mL vials (Labco exetainers, UK), avoiding transfer of liquid. Room air samples (20 mL) were collected at three timepoints (start, middle, end) to allow for baseline determination.

Samples from three to four peepers were used for the determination of dissolved ions (nitrate, nitrite, ammonium, Fe(II)/Fe(III), sulfate, sulfide), porewater pH and electric conductivity as well as organic acids, and one peeper was used for the determination of dissolved carbon fractions (organic, inorganic, total). Samples from one peeper were analyzed for dissolved organic carbon (DOC) using a Shimadzu TOC-L analyzer (Shimadzu, Japan). All samples were filtered through 0.2 µm PES filters prior to analyses.

### Enumeration of functional microbial groups by most probable number (MPN) counts

Most probable number counts were set up to target aerobic and anaerobic heterotrophs, nitrifiers, denitrifiers, Fe(III) reducers and sulfate reducers in peat soil from 0 to 20 cm (surface) and 40 to 60 cm (subsurface). Serial dilutions of both peat layers ranging from 10^-2^ to 10^-8^ were prepared in sterile 1x PBS. All growth media were prepared in artificial porewater (as described in (Kujala *et al*. 2020)) at a final pH of 7 and supplemented as follows: 1.25 g L^-1^ nutrient broth (aerobic and anaerobic heterotrophs), 5 mM ammonium and 5 mM bicarbonate (nitrifiers), 5 mM nitrate, 5 mM lactate and 5 mM acetate (denitrifiers), 1 mM Fe(III), 5 mM lactate and 5 mM acetate (Fe(III) reducers), and 5 mM sulfate, 5 mM lactate, 5 mM acetate (sulfate reducers). 225 µl of the respective growth medium were inoculated with 25 µl diluted sample in 96-well plates in 6 replicates per dilution step and peat layer. Negative control wells were inoculated with sterile PBS in 12 replicates per plate. MPN incubations targeting aerobic microorganisms (aerobic heterotrophs, nitrifiers) were incubated under ambient air in the dark, while MPN incubations targeting anaerobic microorganisms (anaerobic heterotrophs, denitrifiers, Fe(III) reducers, sulfate reducers) were incubated in air-tight containers under an anoxic atmosphere generated using AnaeroGen pouches (Thermo Fisher Scientific) in the dark. Growth of microorganisms in the plates was monitored by determining optical density (OD) at 600 nm at the start, at the end of week one and after three weeks. In addition, indicators of process activity was assessed after three weeks by determining the presence of nitrate and/or nitrite (nitrifiers and denitrifiers), Fe(II) (Fe(III) reducers) or sulfide (sulfate reducers). Growth and process positive wells per dilution were counted and MPNs calculated for each functional group and peat layer.

### Microcosm incubations

Slurry incubations were set up with field-fresh peat from 0 to 20 cm (surface) and 40 to 60 cm (subsurface) to test the process potential of the following functional groups: i) nitrifiers, ii) denitrifiers, iii) Fe(III) reducers, iv) sulfate reducers, v) fermenters and methanogens. Homogenized soil was diluted 1:5 with milliQ H_2_O (10 g soil + 40 mL ddH_2_O) in 100 mL serum bottles in three replicates per depth and functional group. To test the nitrification potential, bottle openings were covered with pierced parafilm to allow air exchange in the bottles. To test anaerobic process potentials, bottles were crimp sealed with butyl rubber stoppers and the headspace was flushed with N_2_ to establish an anoxic atmosphere. Electron donors/acceptors were supplemented as follows: i) 200 µM ammonium to assess the peat’s nitrification potential, ii) 200 µM nitrate to assess the peat’s denitrification potential, iii) 200 µM Fe(III) to assess the peat’s Fe(III) reduction potential, iv) 200 µM sulfate to assess the peat’s sulfate reduction potential, and v) 1000 µM N-acetylglucosamine (NAG) to assess the peat’s fermentation and methanogenesis potential. Slurry incubations were sampled daily using a syringe (withdrawing ∼1 mL of liquid) during the first week of incubation and at less frequent intervals for the following three weeks. Samples were filtered (0.2 µm PES filter) and stored frozen prior to further analysis, unless the analyte was unstable and required immediate analysis (Fe(II), sulfide). The headspace of the incubations targeting fermenters and methanogens was sampled on days 0, 2, 4 and 11. 1 mL headspace gas was withdrawn with a syringe and injected into 15 mL vials (Labco exetainers, UK) that had been prefilled with 19 mL N_2_ to achieve a 1:20 dilution.

### DNA extraction and amplicon sequencing of bacterial and archaeal communities

Peat samples were thawed, and DNA was extracted using the ZymoBiomics DNA Miniprep Kit according to the manufacturer’s instructions. Bacterial and archaeal full-length 16S rRNA genes were amplified using the primer pairs 27F (5’-AGA GTT TGA TCM TGG CTC AG-3’) + 1492R (5’-GGT TAC CTT GTT ACG ACT T-3’) (Lane 1991) and 8F (5’-TCC GGT TGA TCC TGC C-3’; (Teske *et al*. 2002)) + 1492R, respectively. For full-length sequencing of bacterial and archaeal 16S amplicons, PCR products were barcoded using the Oxford Nanopore Technology (ONT) Native Barcoding kit V14 with 24 barcodes (SQK-NBD114.24; one library each for archaeal and bacterial amplicons) and sequenced on a MinION R10.4.1 flow cell according to the manufacturer’s instructions. Basecalling and demultiplexing was done with the Dorado basecaller (<u>GitHub - nanoporetech/dorado: Oxford Nanopore’s Basecaller</u>) using the Super accurate basecalling model, discarding sequences with a quality score below 10. Demultiplexed sequence reads were imported into qiime2 version 24.10 (Bolyen *et al*. 2019). For each barcode, forward and reverse primers were initially processed separately due to the mixed orientation of Nanopore reads. Forward/reverse primer were identified and trimmed at the 5’-end of the reads and the reverse complements of the reverse/forward primer were identified and trimmed at the 3’-end of the reads using the q2-cutadapt plugin (Martin 2011). Reads in which primers were not detected at both ends were discarded. The sequence orientation of the reverse reads was adjusted using the q2-rescript plugin (Robeson *et al*. 2021), after which forward and reverse reads and OTU tables were merged. Sequences were classified against the greengenes2 database (McDonald *et al*. 2024) using the consensus blast q2-feature-classifier (Camacho *et al*. 2009). Unassigned reads and reads with taxonomy assigned only at domain-level were removed from the OTU table. The OTU table was collapsed at species-level and rarefied to an even sampling depth of 1000 sequences. Alpha (Shannon diversity (Shannon 1948) and Pielou Evenness (Pielou 1966)) and beta diversity (Bray and Curtis 1957) parameters were calculated based on the rarefied OTU table using the q2-diversity plugin.

### Analytical procedures

Gravimetric moisture content of peat samples was determined by comparing the weight of field-fresh peat to the weight of peat dried at 30 °C peat for 7 days. Water extracts were prepared by diluting field-fresh peat soil 1:5 with ddH_2_O, shaking at 350 rpm for 1 hour on a rotary shaker and filtering through a 0.2 µm filter. Concentrations of analytes in samples from microcosm incubations, in porewater extracted from peepers and in peat extracts were determined colorimetrically using established protocols for ammonium (Fawcett and Scott 1960), nitrate (Miranda, Espey and Wink 2001), nitrite (Miranda, Espey and Wink 2001), sulfate (Dodgson 1961), sulfide (Cline 1969), total iron, and Fe(II) (Fortune and Mellon 1938). Concentrations of N_2_O, CO_2_ and CH_4_ from porewater or microcosm samples were determined using a gas chromatograph (8890 GC System, Agilent, USA) at the Natural Resources Institute Finland, Oulu laboratory. Organic acids were quantified from 0.2 µm filtered water samples taken from extracted porewater and the fermenter/methanogen microcosms using an Agilent 1200 HPLC system equipped with an Aminex HPX-87H column (300 x 7.8 mm; Biorad) and a UV-visible light diode array detector (detection at 210 nm) using a mobile phase of 4 mM H_3_PO_4_, a column temperature of 60°C and a flow rate of 0.6 mL/min. Organic acid standards with known concentrations were used for calibration. Total elemental concentrations in peat were determined from pooled samples after microwave digestion via ICP-OES analysis at Eurofins Ahma Oy (EPA 3051A:2007).

## RESULTS AND DISCUSSION

### High microbial diversity and abundance of functional groups in Puukkosuo fen soil

Archaeal and bacterial communities were assessed through sequencing of full-length 16S rRNA gene amplicons. Communities changed with depth, and the highest diversity (no. of observed species, Shannon diversity and Pielou Evenness) was observed in the top peat layers (0 to 30 cm; Fig. S1). Both the archaeal and bacterial communities in the upper 10 cm differed strongly from the communities in all other layers, with less pronounced differences in community composition between the lower layers (Fig. S2). Archaeal communities were dominated by sequences affiliated with the *Thermoproteota* genus Ca. *Bathycorpusculum*, the *Halobacteriota* genera *Methanothrix,* Ca. *Methanoflorens* and the *Thermoplasmata* family *Methanomassiliicoccaeae* (Fig. 2A). The relative abundance of Ca. *Bathycorpusculum* affiliated sequences increased with increasing depth from 22 to 63% while the relative abundances of *Methanothrix* and Ca. *Methanoflorens* affiliated sequences decreased. The phylum *Bathyarchaeota* is widespread in anoxic sediments and is known for its heterotrophic pathways of energy conservation (Zhou *et al*. 2018b). Genomic studies have indicated that members of this phylum encode pathways for the potential to anaerobically utilize a variety of organic compounds, including proteins, carbohydrates, fatty acids, methane, and other organic matter, as well as other metabolic capabilities such as acetogenesis or dissimilatory nitrogen and sulfur reduction (Zhou *et al*. 2018b; Hou *et al*. 2023). MAGs of the candidate genus *Bathycorpusculum* have the genetic potential for fermentative metabolism and can potentially use H_2_ and amino acids as electron donors for reductive acetogenesis from CO_2_ or even methylated compounds (Loh, Hervé and Brune 2021; Protasov *et al*. 2023). However, unlike some other *Bathyarchaeota*, *Bathycorpusculum* does not encode methyl-coenzyme M reductase and is thus likely not methanogenic (Loh, Hervé and Brune 2021). In Puukkosuo, *Bathycorpusculum*-affiliated Archaea might thus play an important role in the breakdown of complex peat organic matter and provide substrates for subsequent fermentations or anaerobic respiration processes by other microorganisms in the community. Of the detected *Halobacteriota*, *Methanoregulaceae* and *Methanoculleaceae* are hydrogenotrophic methanogens (Bapteste, Brochier and Boucher 2005), while *Methanothrix* are known for aceticlastic methanogenesis. Ca. *Methanoflorens* likely also is hydrogenotrophic (Mondav *et al*. 2014; Woodcroft *et al*. 2018; Bräuer *et al*. 2020). Members of the *Methanomassiliicoccaceae* are considered methylotrophic methanogens and are widely distributed in wetland soils (Söllinger *et al*. 2016; Narrowe *et al*. 2019; Bräuer *et al*. 2020). Thus, hydrogenotrophic, aceticlastic and methylotrophic methanogenesis are all feasible in Puukkosuo fen, with hydrogenotrophic, aceticlastic and methylotrophic taxa accounting for approximately 61, 31 and 8% of the methanogenic community. Indeed, methanogenesis in Puukkosuo fen is stimulated by H_2_/CO_2_, formate, acetate, and methanol, with H_2_/CO_2_ and formate showing the strongest stimulatory effect (Kujala, Schmidt and Horn 2024). The importance of the methylotrophic pathway is only insufficiently studied even though methylotrophic methanogens are detected in a wide range of environments (Conrad 2020). In peatlands, methylotrophic methanogenesis is believed to be of minor importance (Drake, Horn and Wüst 2009), even though methanol released from living or dead plant material is likely present in peat (Galbally and Kirstine 2002). It has been suggested that lack of H_2_ and formate may restrict methylotrophic methane production even if sufficient levels of methanol are available (Bräuer *et al*. 2020). Thus, aceticlastic and especially hydrogenotrophic methanogenesis are likely also the dominant forms of methanogenesis in Puukkosuo fen.

**Figure 2:**
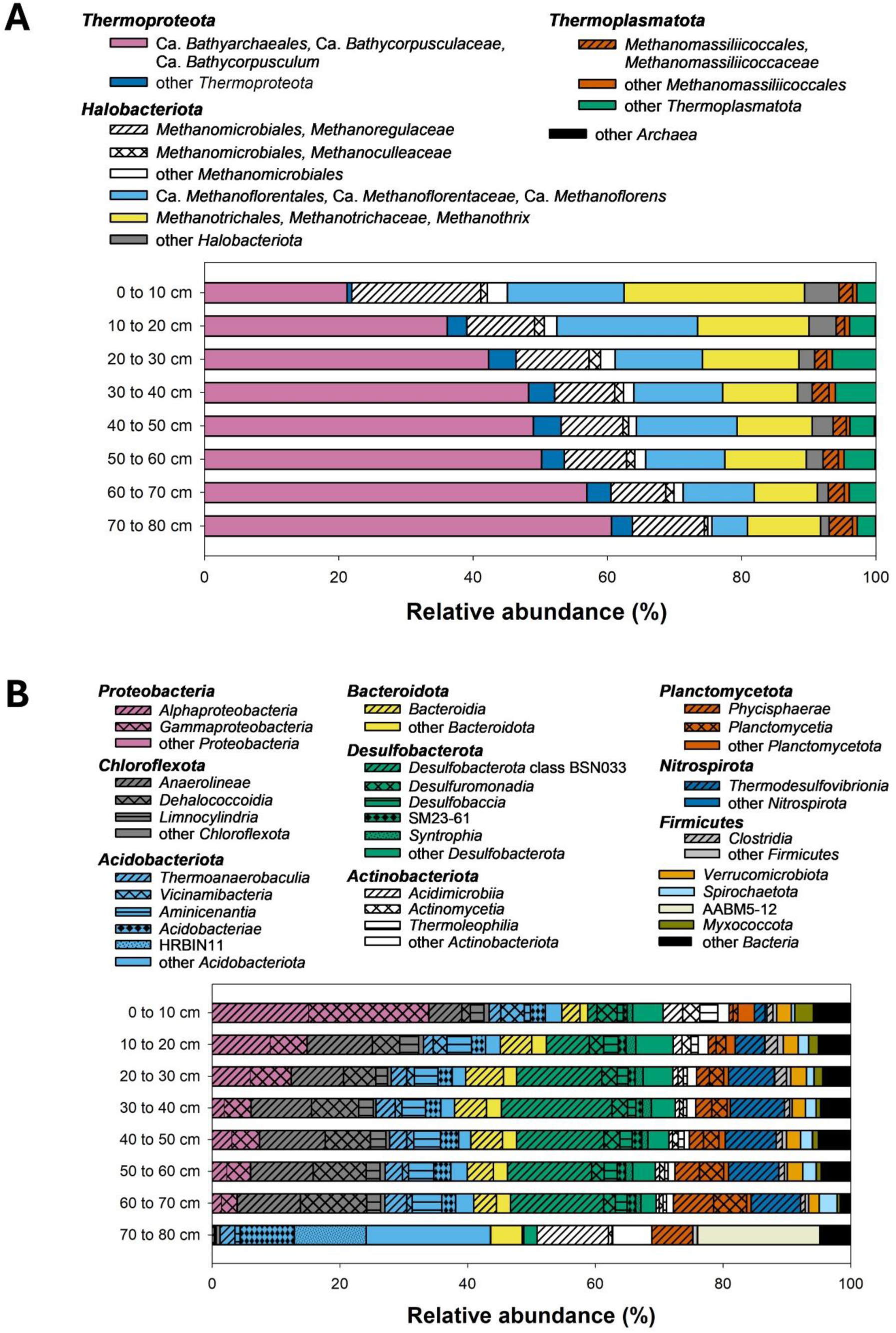
Depth-resolved composition of the archaeal (A) and bacterial (B) communities in Puukkosuo fen. Community composition is based on sequencing of full-length archaeal and bacterial 16S rRNA gene amplicons obtained from three replicate peat cores (sampled on September 19^th^, 2023). Sequences were taxonomically assigned against the greengenes 2 database, and family- to genus-level (A) or class-level (B) average relative abundances of the three replicates are displayed. Classes/families/genera with an average relative abundance <1% were grouped on the next higher taxonomic level. Phyla with an average relative abundance <1% are grouped as “other Archaea/Bacteria”.

Bacterial communities were considerably more diverse than archaeal communities with approximately 10x higher numbers of detected species (Fig. S1), and were dominated by sequences affiliated with *Proteobacteria, Chloroflexota, Acidobacteriota, Bacteroidota* and *Desulfobacterota* (Fig. 2B). While *Proteobacteria* accounted for >30% of sequence in 0 to 10 cm peat, their abundance decreased steadily to below <1% in 70 to 80 cm peat. In 70 to 80 cm peat, the bacterial communities were dominated by sequences affiliated with *Acidobacteriota, Actinobacteriota* and AABM5-12 (Fig. 2B). Sequences were affiliated with taxa capable of anaerobic metabolism such as fermentation or anaerobic respirations like denitrification, iron reduction and sulfate reduction, as well as aerobic nitrifiers and methanotrophs. Primary fermenters (e.g., mixed acid fermentation in *Clostridia*) or secondary (syntrophic) fermenters (e.g., propionate-utilizing members within the *Desulfobacterota*) play an essential role in the stepwise breakdown of organic carbon compounds and provide substrates for anaerobic respiration (Drake, Horn and Wüst 2009; Morris *et al*. 2013; Westerholm, Calusinska and Dolfing 2022). Some of the syntrophic *Desulfobacterota* (e.g., *Syntrophobacter*) can also grow independently as sulfate reducers (Westerholm, Calusinska and Dolfing 2022). Sulfate reducers are important for ecosystem functioning and carbon turnover, e.g., in wetlands (Demin et al., 2024). Potentially sulfate-reducing bacteria are e.g., found in the phyla *Desulfobacterota* and *Acidobacteriota* (Hausmann *et al*. 2018; Vigneron *et al*. 2018; Dyksma and Pester 2023), which were both abundant in Puukkosuo fen soil (Fig. 2B). Sulfate reducing bacteria often have adaptive capabilities that allow them to adjust to changes in environmental conditions, and some can degrade complex polysaccharides such as pectin (Hausmann *et al*. 2018; Dyksma and Pester 2023). Iron reducers and denitrifiers likewise belong to many different taxa, and sequences from Puukkosuo fen were affiliated with potentially iron-reducing and denitrifying genera such as *Geobacter, Rhodoferax* or *Acidithiobacillus* and *Bradyrhizobium, Anaeromyxobacter* or *Accumulibacter*. *Geobacter* and *Rhodoferax* reduce iron at circumneutral pH (Lovley *et al*. 1993; Kato and Ohkuma 2021) and might thus be involved in iron reduction in Puukkosuo fen. Ongoing iron reduction has been associated with increased degradation of peat organic matter and DOC release (Küsel *et al*. 2008; Pan *et al*. 2016; Chen *et al*. 2020), and iron reducers in Puukkosuo fen might thus accelerate organic matter degradation. Sequences were also affiliated with known methanotrophic taxa within the *Methylococcales* (e.g., *Methylococcus, Methylomonas*)*, Rhizobiales* (e.g., *Methylosinus, Methylocystis*) and *Verrucomicrobiota* (*Methylacidiohilaceae*), and their combined relative abundance ranged from 3.4% in 0 to 10 cm to 0.4% in 70 to 80 cm. Relative abundances of methanotrophs in surface peat have been reported in the range from 2.6-5.3% (Nweze *et al*. 2024). Bacterial nitrifiers such as *Nitrosospira, Nitrosococcus* or *Nitrospira* had a low relative abundance (0.08-0.35%) which was likewise highest in 0 to 10 cm peat. Aerobic methanotrophy and nitrification are thus likely restricted to the upper soil layers in Puukkosuo fen.

Enumeration of functional microbial groups through MPN counts in surface (0 to 20 cm) and deeper (40 to 60 cm) peat revealed aerobic heterotrophs, anaerobic heterotrophs and sulfate reducers to be the most abundant functional groups with ≥ 10^6^ * g_DW_^-1^ cultivable cells in both layers (Fig. 3). Total microbial cell counts in peat soil estimated through qPCR or microscopy are typically in the order of 10^9^ to 10^11^ cells * g_DW_^-1^ (Dedysh *et al*. 2006; Reiche, Torburg and Küsel 2008; Palmer, Drake and Horn 2010; Steger *et al*. 2011; Prasitwuttisak *et al*. 2022), and abundance of culturable aerobic and anaerobic heterotrophs can range from 10^7^ to 10^9^ cells * g_DW_^-1^ (Wüst, Horn and Drake 2009; Kujala *et al*. 2020; Heitkämper *et al*. 2024). Abundance of sulfate reducers on the other hand are typically slightly lower (Castro, Reddy and Ogram 2002; He *et al*. 2010; Steger *et al*. 2011), and they have even been considered ‘rare biosphere’ in some systems (Demin *et al*. 2024). Nitrifiers, nitrate reducers, denitrifiers and iron reducers had slightly higher numbers of cultivable cells in upper peat than in lower peat, with mean cell number of 2.5 * 10^4^, 1.4 * 10^6^, 8.1 * 10^4^, and 1.4 * 10^5^ g_DW_^-1^ cultivable cells, respectively. This is in good agreement with previous studies which have likewise shown highest numbers in surface peat and reported abundances in a similar range (Reiche, Torburg and Küsel 2008; Wüst, Horn and Drake 2009; Palmer, Drake and Horn 2010; Naylor, McClure and Jansson 2022). Low numbers of culturable nitrifiers also reflect the low relative abundance of nitrifying genera in the microbial communities. Nitrifier abundance and activity in water-logged soils is often low, as low oxygen availability limits this very specialized functional group (Nguyen *et al*. 2018; Zhou *et al*. 2018a).

**Figure 3:**
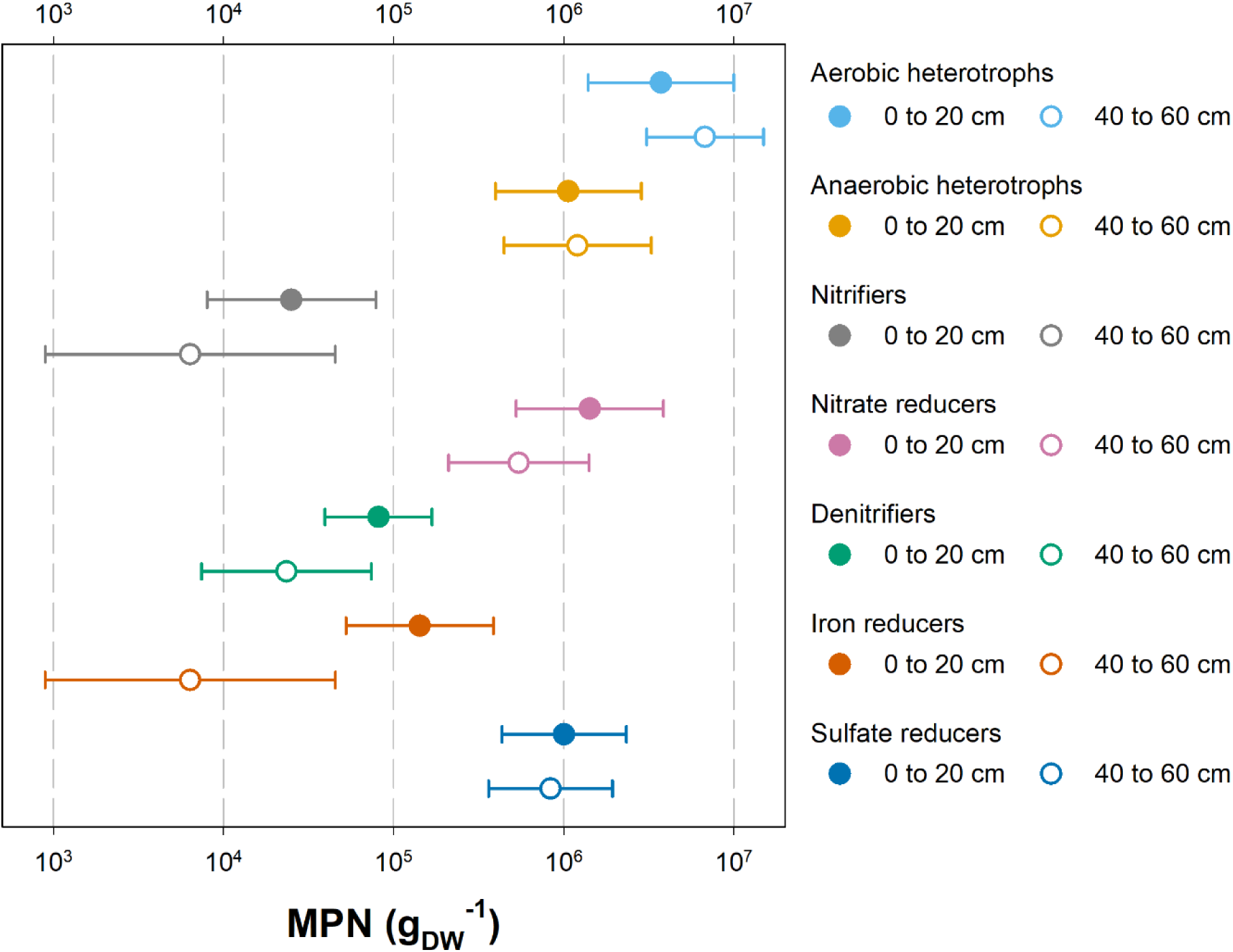
Abundance of different aerobic and anaerobic functional groups in surface (0 to 20 cm) and subsurface (40 to 60 cm) peat from Puukkosuo fen (sampled September 14th, 2023). Most probable number (MPN) counts of aerobic and anaerobic heterotrophs, nitrifiers, nitrate reducers, denitrifiers, iron reducers, and sulfate reducers were determined through serial dilutions of field-fresh peat soil (10^-2^ to 10^-8^) inoculated in six replicates into growth medium specific for the respective functional group and incubated for three weeks in the dark under oxic or anoxic conditions.

### Functional potential of Puukkosuo fen microbes

Peatland microbes have the potential for versatile metabolic functions that contribute to the predominantly anaerobic degradation of organic matter, as aerobic processes are limited to surface layers where oxygen is available. Microcosm incubations with field-fresh peat soil from Puukkosuo fen demonstrated process potential for nitrification, denitrification, iron and sulfur reduction as well as fermentation and methanogenesis (Fig. 4, 5). Potential nitrification activities were low. In microcosms with surface peat supplemented with 100 µM ammonium, ammonium concentrations decreased slightly over 6 days, while in microcosms with subsurface peat a slight increase in ammonium concentration was observed (Fig. 4A). No net formation of nitrate or nitrite was observed in microcosms with peat from either layer, indicating that ammonium and nitrite oxidation rates were either very low or produced nitrite or nitrate were immediately consumed. The prevalence of anoxic conditions throughout most of the peat likely does not favor the establishment of nitrifier communities or strong nitrification potentials. In microcosms set up to determine denitrification potential, nitrate concentrations decreased in both layers without apparent delay (Fig. 4B). Consumption of 100 µM supplemented nitrate was slightly faster in surface peat, but >90% of supplemented nitrate was consumed in both layers after 3 days of incubation. Nitrite as an intermediate in the denitrification process transiently accumulated in the first 2 days of incubation, with higher accumulation in microcosms with subsurface peat, but was completely depleted in microcosms with soil from both depths after 4 days of incubation, indicating its further reduction to nitrogen gases. Previous studies conducted in Puukkosuo fen as well as in other peatlands have likewise demonstrated high denitrification activities especially in surface layer peat and production of N_2_O or N_2_ as denitrification end products (Van Beek *et al*. 2004; Palmer, Drake and Horn 2010; Palmer and Horn 2015). Ammonium accumulated in the microcosms after 2-3 days (∼40 and 20 µM in microcosms with surface and subsurface peat soil, respectively), which could also hint at dissimilatory nitrate reduction to ammonium (DNRA). The contribution of DNRA to nitrate turnover in peatlands is variable and influenced by many factors(Rütting *et al*. 2011; Shi *et al*. 2021). Redox status and C/N ratio have been proposed as potential factors determining the contribution of DNRA, whereby higher carbon and oxygen availability favors ammonium production (Rütting *et al*. 2011). However, ammonium was also extractable from peat in similar concentrations to those measured in the microcosms (Table S1), and detected ammonium might thus (in part) be due to ammonium leaching from peat rather than its production via DNRA. Iron reduction potential was observed in microcosms with peat from both depths, with higher dissolved Fe(II) concentrations produced in microcosms with surface peat (Fig. 4C). Fe(II) production rates were low in comparison to those reported from other peatlands (Reiche, Torburg and Küsel 2008). However, in the present study only dissolved Fe(II) was determined, which would omit any Fe(II) bound to the peat which might e.g., have been HCl-extractable. Sulfate reduction potential was likewise higher in microcosms with surface peat, with >95% of supplemented sulfate being consumed after 7 days, while in microcosms with subsurface peat 80% of supplemented sulfate were consumed after 26 days (Fig. 4D). Sulfide formation was observed in microcosms with peat from both depths. In microcosms with surface peat, only 25% of the consumed sulfate were recovered as sulfide, while in microcosms with subsurface peat sulfate consumption equaled sulfide production. The mismatch between sulfate consumption and sulfide production might be explained adsorption or incorporation of sulfide into the peat substance or precipitation of sulfide-iron minerals (Bottrell and Coulson 2003; Wang *et al*. 2025). Fermentation and methanogenesis potentials were determined in microcosms supplemented with 1000 µM NAG. NAG is a monomer of chitin as well as of bacterial cell wall peptidoglycan, and thus likely constitutes an important substrate for microorganisms in peatlands (Tveit *et al*. 2015; Pazos and Peters 2019; Wörner and Pester 2019). NAG concentrations decreased steadily in microcosms with peat soil from both layers, and NAG was completely consumed after 10 days of incubation (Fig. 5A). Acetate accumulated to concentrations of approximately 2500 µM in both layers by day 10, while other organic acids were detected in much lower concentrations. Earlier studies in peatlands (including Puukkosuo fen) likewise demonstrated high acetate production potential (Hädrich *et al*. 2012; Ye *et al*. 2014; Kujala, Schmidt and Horn 2024). Acetate turnover in peat is complex, with different processes contributing to its production and consumption. Acetate can be formed by fermentation or by acetogenesis and is a substrate in the aceticlastic methanogenesis, but also as electron donor for anaerobic respirations like denitrification or sulfate reduction (Drake, Horn and Wüst 2009; Hädrich *et al*. 2012; Ye *et al*. 2014). Low amounts of butyrate and propionate were detected at the end of the incubation (∼75 µM and ∼150 µM, respectively) in both layers, while formate transiently accumulated to 195 µM (max. at day 4) in microcosms with lower layer peat (Fig. 5A). Formate, butyrate and propionate fuel methanogenesis either directly (formate) or indirectly (butyrate, propionate) through syntrophic interactions with secondary fermenters (Drake, Horn and Wüst 2009). Both peat layers produced nearly equal amounts of CO_2_ over the 11-day incubation period (Fig. 5B), while CH_4_ production was approximately double in microcosms with surface peat (Fig. 5C). A decline in CH_4_ production potential with depth has also been described in other peatlands (Williams and Crawford 1984; Küsel *et al*. 2008; Wüst, Horn and Drake 2009).

**Figure 4:**
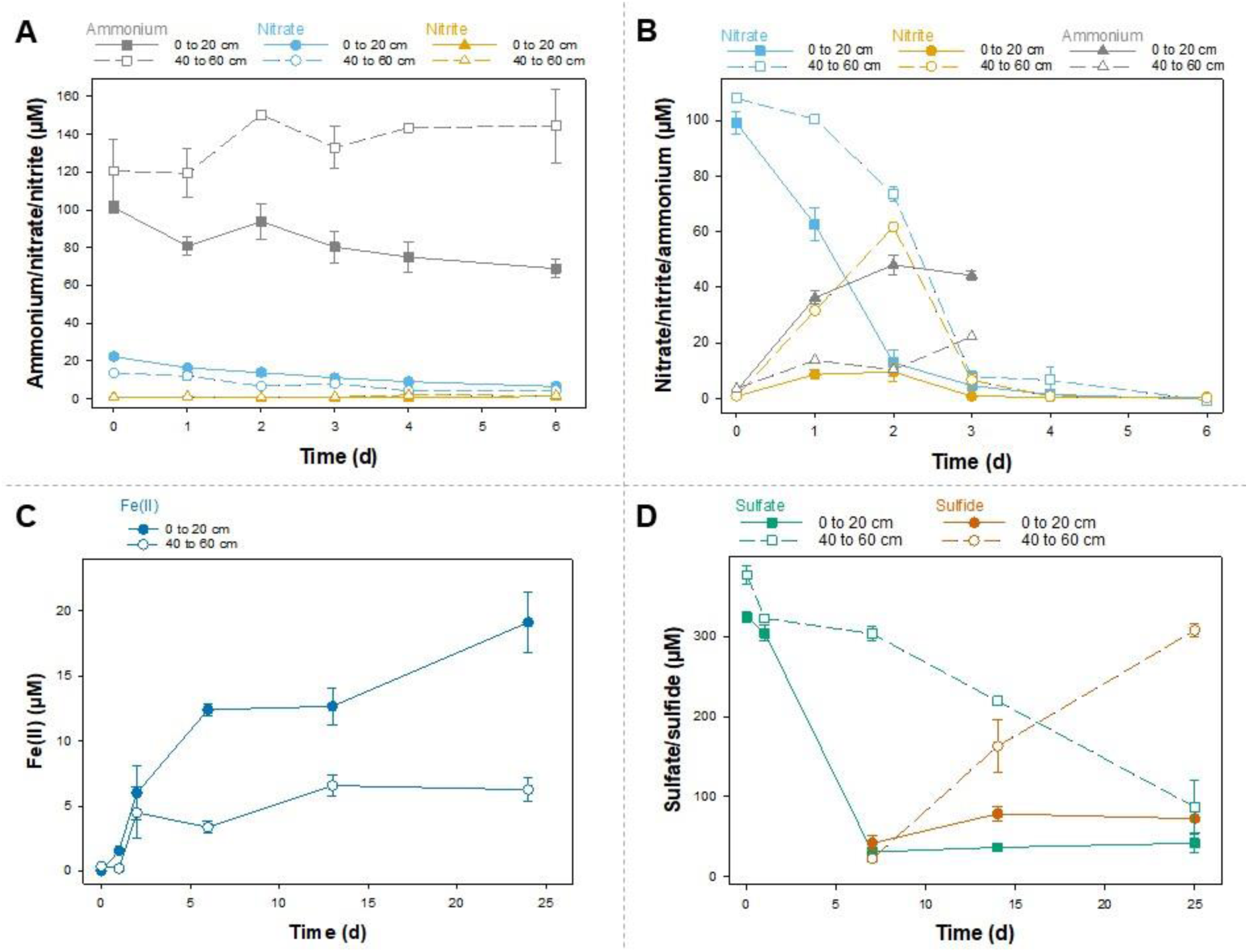
Impact of depth on nitrification (A), denitrification (B), iron reduction (C), and sulfate reduction (D) in microcosms with peat from Puukkosuo fen (sampled September 14th, 2023). Microcosms were prepared in triplicate as 1:5 watery dilutions of surface (0 to 20 cm) and subsurface (20 to 40 cm) peat and incubated for up to 25 days under oxic (A) or anoxic conditions (B-D). Microcosms were supplemented with 100 µM ammonium (A), 100 µM nitrate (B), 100 µM iron (C), and 300 µM sulfate (D) and incubated at room temperature in the dark. The liquid phase was sampled in regular intervals, and concentrations of compounds relevant to the respective process were determined.

**Figure 5:**
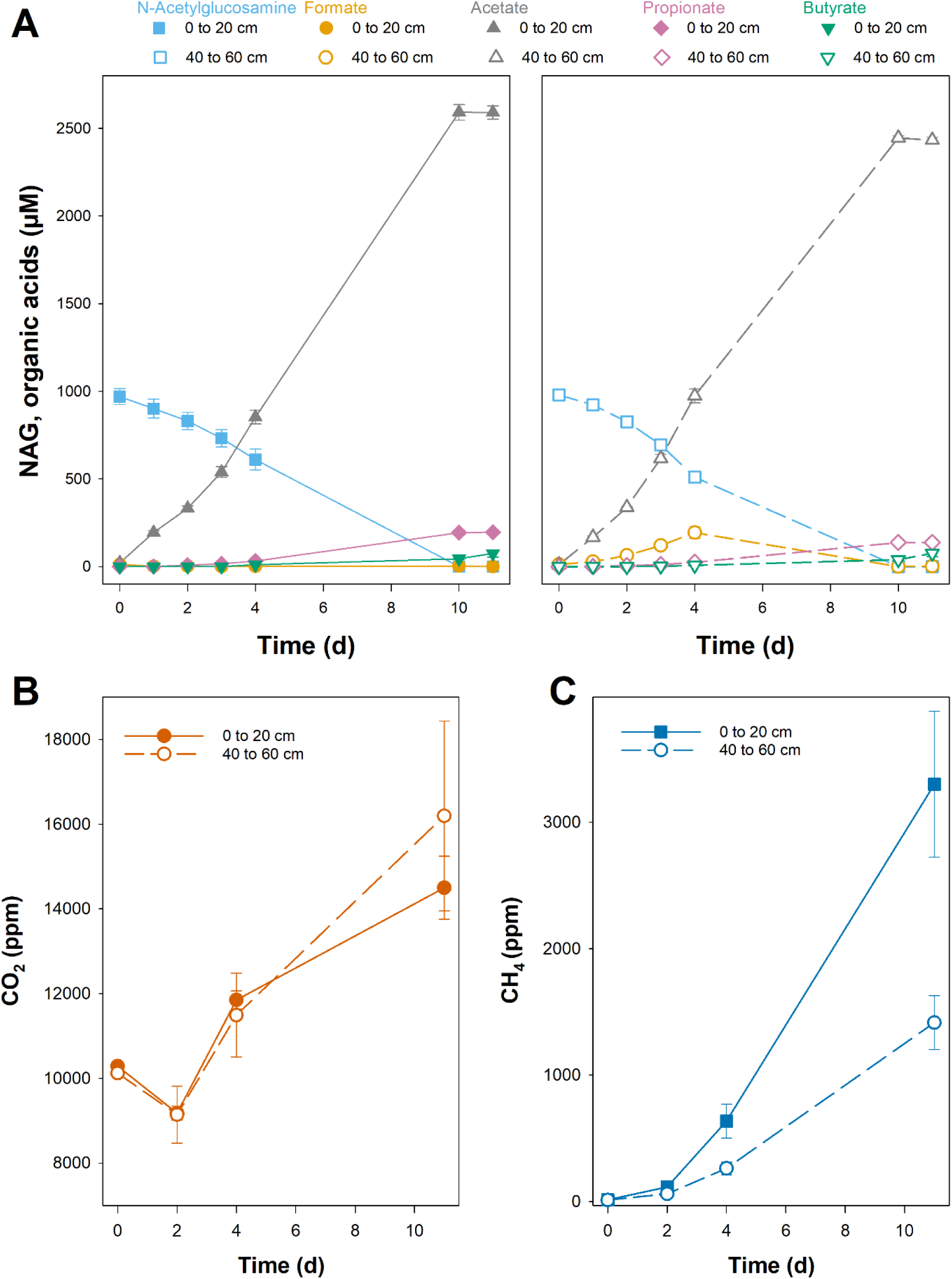
Impact of depth on fermentation and methanogenesis in microcosms with peat from Puukkosuo fen (sampled September 14th, 2023). Microcosms were prepared in triplicate as 1:5 watery dilutions of surface (0 to 20 cm) and subsurface (20 to 40 cm) peat and incubated for 11 days under anoxic conditions. Microcosms were supplemented with 1000 µM N-acetylglucosamine (NAG) and incubated at room temperature in the dark. The liquid (A) and gas (B, C) was sampled in regular intervals, and concentrations of NAG, organic acids, CO2 and CH4 were determined.

### Combined action: In situ concentrations in different depth of Puukkosuo fen

Puukkosuo fen can serve as a model ecosystem to study aerobic and anaerobic processes involved in organic matter degradation and turnover as well as their interactions. These processes occur in part in parallel but also in different layers. Process rates are governed by environmental conditions (oxygen, pH), substrate and electron acceptor availability as well as microbial community composition and activity. While laboratory incubations provide a good opportunity to assess the process potential of specific functional groups, they can only insufficiently address the interplay of different processes and microbial groups in the peat column. Insight can be gained by studying the concentrations of process substrates, products and intermediates in solids, liquids, and gases to search for traces of microbial *in situ* activity. The porewater pH in Puukkosuo fen was neutral, ranging from 6.9 to 7.2 with slightly higher values in the lower peat layers. Thus, ongoing processes in Puukkosuo fen might differ from those described in other peatlands, in which the pH is often more acidic (Goodwin and Zeikus 1987; Blodau *et al*. 2007; Küsel *et al*. 2008; Wüst, Horn and Drake 2009; Frank *et al*. 2014).

The surface layer of peatlands often undergo watertable fluctuations related e.g., to dry periods, precipitation events or spring snowmelt which allow for intrusion of oxygen (Albert-Saiz *et al*. 2025). However, oxygen concentrations decline sharply with depth, even when peat is not water-saturated (Dickopp, Lengerer and Kazda 2018). In Puukkosuo fen, peat was highly water saturated with a water content ranging from 87 to 92% with a slightly declining trend towards in lower layers (Table S1). Oxygen penetration into the peat porewater was restricted to the first 10 cm, below which oxygen was not detected anymore (Fig. 6), indicating that in peat below 10 cm anaerobic processes increase in importance and degradation processes are governed by the availability of alternative electron acceptors. Presence of alternative electron acceptors in high concentrations can suppress methanogenesis (Knorr, Lischeid and Blodau 2009; Kujala, Schmidt and Horn 2024). However, concentrations of alternative electron acceptors such as nitrate, Fe(III) and sulfate in porewater were low throughout the porewater profile with concentrations ranging from 4 to 6 µM, 0 to 2 µM and 5 to 20 µM, respectively (Fig. 6). Concentrations of alternative electron acceptors in peat porewater are highly spatially and temporarily variable (Blodau *et al*. 2007; Schmalenberger, Drake and Küsel 2007; Küsel *et al*. 2008; Knorr, Lischeid and Blodau 2009; Maljanen *et al*. 2010; Frank *et al*. 2014). Even though only low concentrations are detected in peat or porewater, turnover of said electron acceptors might still be ongoing, which may in fact be the reason for the low concentrations (Palmer, Drake and Horn 2010; Pester *et al*. 2012), or electron acceptors such as Fe(III) might be bound to peat rather than being dissolved in porewater, and repeated measurements of porewater concentrations in Puukkosuo fen would give valuable insights into seasonal electron acceptor availability in different peat depths. Internal cycling of Fe or sulfate could help to replenish Fe(III) and sulfate (Blodau *et al*. 2007; Pester *et al*. 2012; Patzner *et al*. 2022). Reoxidation of these electron acceptors typically occurs at or just below the oxic-anoxic interface (Brune, Frenzel and Cypionka 2000; Blodau *et al*. 2007). Looking at potential products or intermediates in the denitrification, Fe(III) reduction and sulfate reduction processes, N_2_O as an intermediate of denitrification was observed in low concentrations and showed a declining trend from the surface (117 nM) to 80 cm depth (54 nM), indicating highest N_2_O net production near the surface (Fig. 6). Fe(II) and sulfide were mainly detected in 20 to 40 cm depth, and total peat sulfur concentrations were likewise high in those depths (Fig. 6). Sulfur might be immobilized in the peat as Fe(II)-sulfide after reduction of both Fe(III) and sulfate (Bottrell and Coulson 2003; Pester *et al*. 2012; Wang *et al*. 2025). In addition to Fe and sulfur, small concentrations of the metals manganese, cobalt, copper, molybdenum, nickel, lead and zinc were detected in peat soil, with highest concentrations typically in surface peat (Table S1). Even though some of the detected metals like manganese or copper are also redox-active, they were not investigated further in the present study.

**Figure 6:**
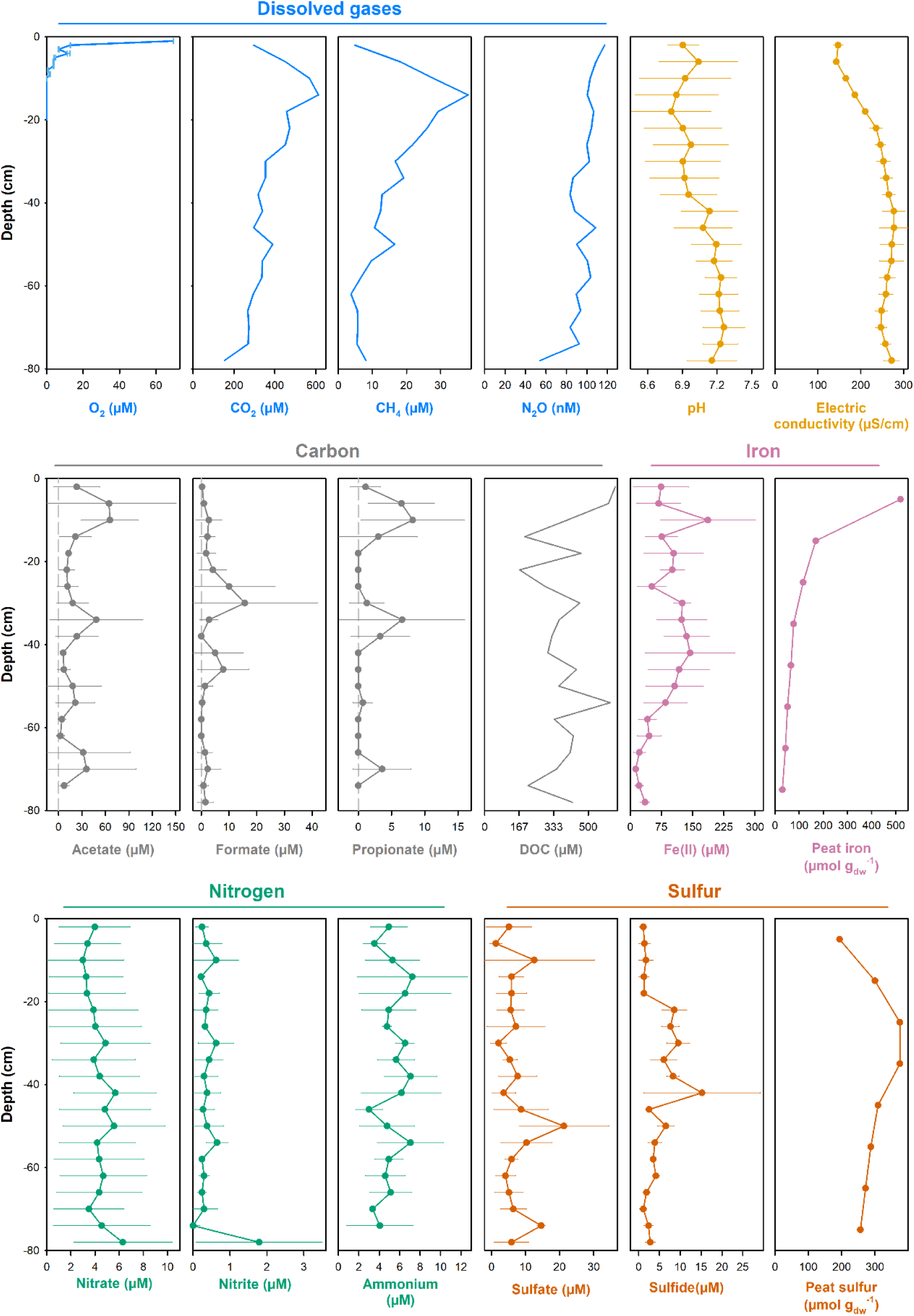
Depth profiles of Puukkosuo peat and porewater. Peat and porewater were sampled on September 19th, 2023) and analyzed for dissolved gases, pH and electric conductivity (EC; top), carbon and iron (middle) as well as nitrogen and sulfur (bottom). For measurements that were conducted on samples from 3 (Fe(II), nitrate, nitrite, ammonium, sulfate, sulfide) or 4 (pH, EC, organic acids) porewater samplers, error bars indicate the standard deviation.

CO_2_, CH_4_ and acetate concentrations peaked at 12 to 16 cm depth with maximum values of 615, 38 µM and 66 µM, respectively, right below the oxic-anoxic interface (Fig. 6). Porewater CH_4_ and acetate profiles in peatlands are highly variable and can differ between sites and seasons, with high CH_4_ concentrations detected in upper as well as in deeper layers (Williams and Crawford 1984; Küsel *et al*. 2008; Knorr, Lischeid and Blodau 2009; Hädrich *et al*. 2012; Sabrekov *et al*. 2024). Net production of CH_4_ has been reported in surface peat near the watertable level, while in deeper layers turnover rates from net production to net consumption of CH_4_ (Knorr, Oosterwoud and Blodau 2008). In Puukkosuo fen, oxygen available in the upper 10 cm might fuel aerobic CH_4_ oxidation, leading to the observed low CH_4_ concentrations in the upper 10-15 cm. Indeed, the oxic-anoxic interface has been considered a hotspot of CH_4_ oxidation, and aerobic methane oxidizers are abundant (Brune, Frenzel and Cypionka 2000; Reis *et al*. 2024). The potential for aerobic methane oxidation in the surface layer of Puukkosuo fen is also supported by the higher relative abundance of methane-oxidizing taxa in surface peat.

### Conclusions, limitations and future perspectives

During a field course at Oulanka research station participants combined forces for the in-depth study of microbial metabolism in pH-neutral Puukkosuo fen. Microbial communities harbored microorganisms with versatile aerobic and anaerobic metabolic capabilities, including potential for nitrification, denitrification, Fe reduction, sulfate reduction, fermentation, and methanogenesis. Measurements of *in situ* concentrations of process substrates, intermediates, or products in peat and porewater indicated the concomitant action of different anaerobic processes in the anoxic peat layers as well as potential methanotrophy in surface peat. Fig. 7 provides a schematic overview of the interplay of processes involved in organic matter degradation in Puukkosuo fen. While the study provides a good insight into various processes feasible in Puukkosuo fen and their potential interplay, it is nonetheless a snapshot: More frequent sampling campaigns would be needed to access the temporal variability in peat and porewater concentrations *in situ* and laboratory studied targeted to specific functional groups (e.g., through isotope labeling experiments) would further help to disentangle the contributions of different microbial groups to overall organic matter turnover in Puukkosuo fen. Despite these limitations, the collected data and the derived interpretations provide valuable steppingstones for future research at the site.

**Figure 7:**
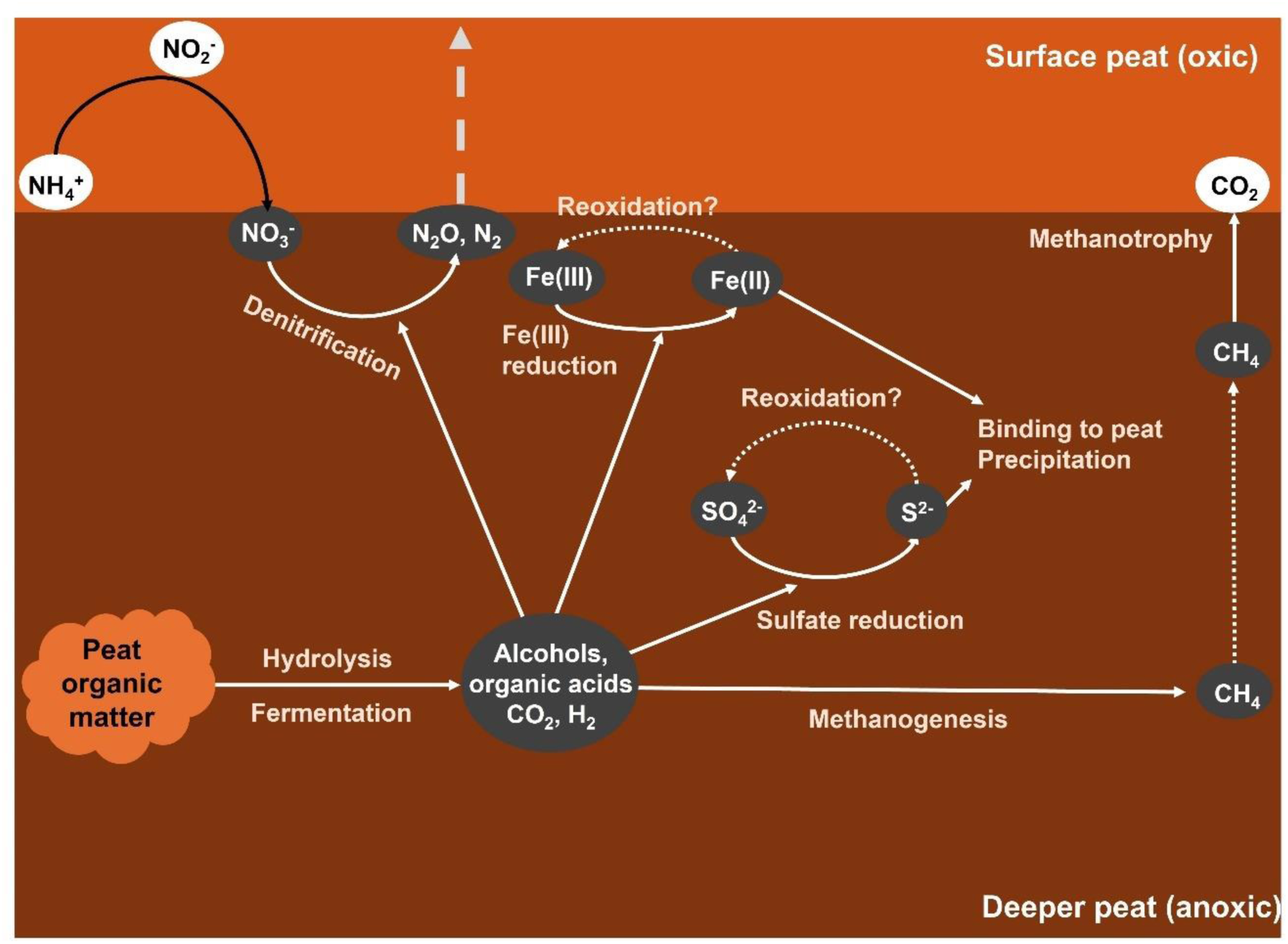
Conceptional model of microbially mediated processes that aid organic matter degradation in Puukkosuo fen. Shown are the major electron donors and acceptors, the processes involved in their turnover and their assumed fate.

## Acknowledgements

The authors thank the team at Oulanka Research station for support during the course (namely Juho Lämsä, Elsi Riihelä, Päivi Salmijärvi, Riku Paavola), Jenni Hultman and Torben Christensen for giving lectures during the course, and the University of Oulu graduate school (UniOGS) for course funding and support in organizational matters.

## Author contributions

The article is based on the results obtained during practical laboratory and field course. Apart from the lead author (KK), who compiled the data and took the main responsibility in writing the manuscript, all other authors have contributed equally and are therefore listed in alphabetical order. KK: Conceptualization, Formal analysis, Investigation, Resources, Supervision, Writing – Original draft, Writing – Review and editing, Visualization, Project administration, Funding acquisition MAH, FPF: Conceptualization, Investigation, Resources, Supervision, Writing – Review and editing VB, JC, SH, EH, IH, PK, EL, JMQ, LP, VP, SS, LT: Formal analysis, Investigation, Writing – Original draft, Writing – Review and editing, Visualization

## Data availability

The sequence data generated in this study has been deposited in the European Nucleotide Archive ENA under study accession number PRJEB102123.

