## Supplemental Figures for "Synergies on many levels: Microbially driven redoxgradients as drivers of greenhouse gas emissions in a pH neutral fen"

### Slide 1
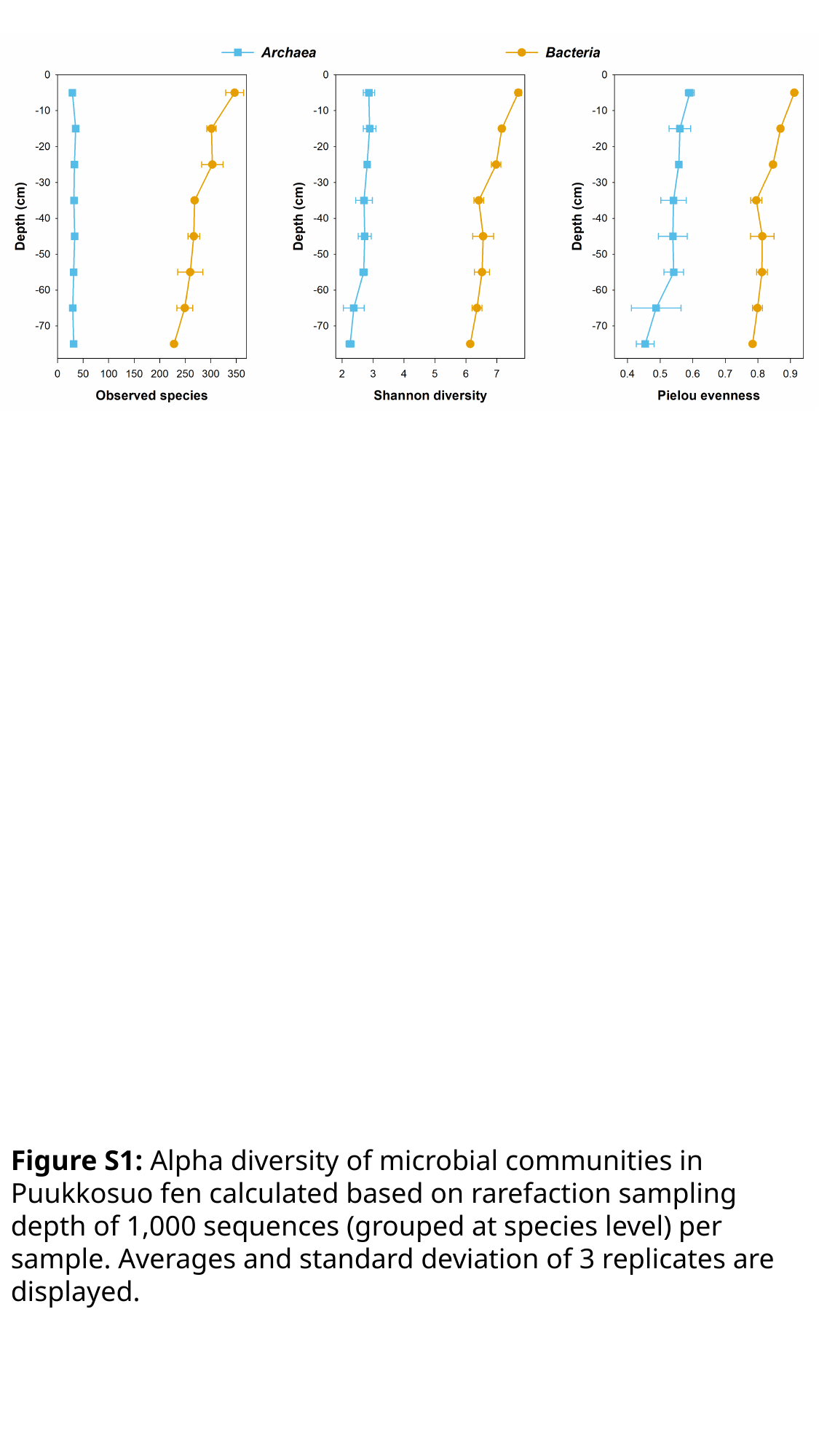

Figure S1: Alpha diversity of microbial communities in Puukkosuo fen calculated based on rarefaction sampling depth of 1,000 sequences (grouped at species level) per sample. Averages and standard deviation of 3 replicates are displayed.

### Slide 2
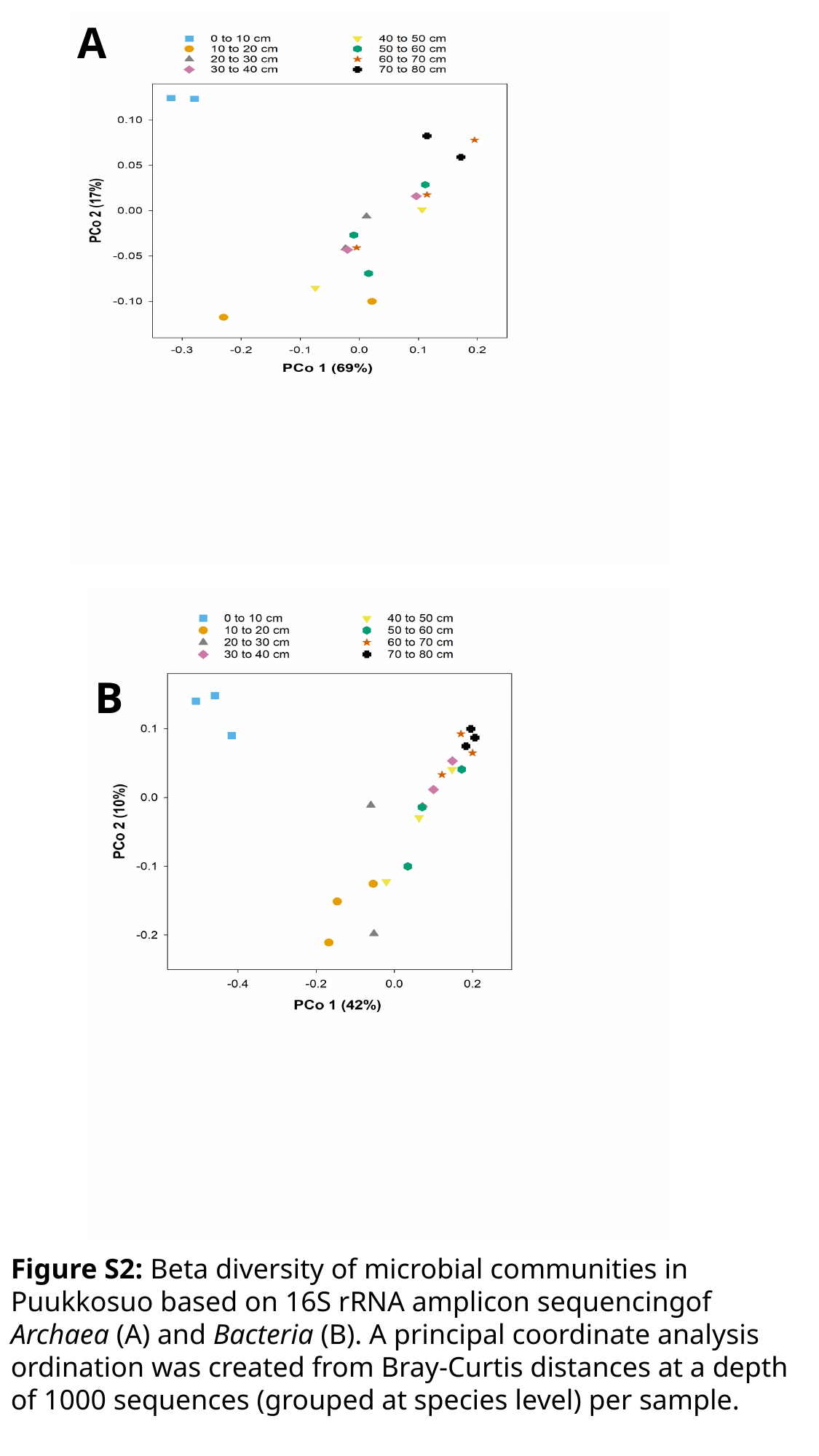

A
B
Figure S2: Beta diversity of microbial communities in Puukkosuo based on 16S rRNA amplicon sequencingof Archaea (A) and Bacteria (B). A principal coordinate analysis ordination was created from Bray-Curtis distances at a depth of 1000 sequences (grouped at species level) per sample.
